## Supplemental Materials for "Genetically Engineered Mice for Combinatorial Cardiovascular Optobiology"

**X-gal staining**

LacZ expression from the optogenetic transgene constructs were screened by X-gal staining (1 mg/mL in 1xPBS; 1mM $MgCl_2$; 250 μM potassium ferricyanide; 250 μM potassium ferrocyanide) at 37ºC for either 4 hrs or overnight. For whole embryo staining, the embryos were fixed in 4% PFA (pH 7.0) for 30 minutes on ice and washed three times in detergent wash buffer (0.1 M phosphate buffer, pH 7.3; 2mM $MgCl_2$; 0.01% Na deoxycholate; 0.02% NP-40) for 20 minutes. Whole tissues were fixed in 4% PFA for 2 hrs on ice, washed three times in 1x PBS for 10 minutes and stained in X-gal. For frozen sections, the fixed tissues were dehydrated in 30% sucrose (1x PBS), embedded in OCT and sectioned at 8 microns. The sections were rinsed in 1x PBS for 5 minutes (3x) and stained in X-gal.

**ImageJ**

The monochrome fluorescent digital images obtained from Leica DMI6000B/Q-imaging or Olympus MVX-10/Andor-iXon CCD cameras were colorized using the Lookup Table command in ImageJ.

**Whole heart clearing**

The neonatal heart was cleared using BABB as described (Joyner & Wall, 2008) and imaged using Olympus OV100 digital camera mounted to a Leica stereoscope.

**Major Resources Table**

### Animals

Information on all lines and the CHROMus project can be found at http://chromus.vet.cornell.edu. CHROMus mice are available from the Jackson Laboratory.

| CHROMus™ ID | Transgenic Line | Jackson Laboratory Stock Number |
|---|---|---|
| B3 | Acta2$^{BAC}$ - RCaMP1.07 | 028345 |
| B4 | αMHC - GCaMP8 | 033341 |
| B5 | αMHC - CatCh2_IRES_lacZ | 030334 |
| B6 | Acta2$^{BAC}$ - Optoα1AR_IRES_lacZ | 028346 |
| B7 | Acta2$^{BAC}$ -Optoß2AR_IRES_lacZ | 028347 |
| B8 | Acta2$^{BAC}$ - CatCh2_IRES_lacZ | 028348 |
| B10 | HCN4$^{BAC}$ -GCaMP8 | 028344 |
| B14 | Acta$^{BAC}$ - GCaMP5_mCherry | 025406 |
| B15 | Cx40$^{BAC}$ - GCaMP5_mCherry | 030333 |
| B16 | Dbh$^{BAC}$ - CatCh2_IRES_lacZ | 032666 |
| B17 | HCN4$^{BAC}$ - CatCh2_IRES_lacZ | 033344 |
| B19 | Acta2$^{BAC}$ - GCaMP2 | 025405 |
| B20 | Cdh5$^{BAC}$ - GCaMP8 | 033342 |
| B22 | Lck$^{BAC}$ - Optoα1AR_IRES_lacZ | 033705 |
| B23 | SP-C$^{BAC}$ - GCaMP8 | 032885 |
| B26 | Cdh5$^{BAC}$ - Optoα1AR_IRES_lacZ | 033343 |
| B27 | Cdh5$^{BAC}$ - Optoß 2AR_IRES_lacZ | 032889 |
| B28 | Cdh5$^{BAC}$ - CatCh2_IRES_lacZ | 033345 |
| B34 | Acta2$^{BAC}$ - GCaMP8.1_mVermilion | 032887 |
| B35 | FoxJ1$^{BAC}$ - GCaMP8.1 | 032888 |
| B36 | Aldh1L1$^{BAC}$ - Optoα1AR_IRES_lacZ | 033706 |

### Figure Supplements

**Table 1 – figure supplement 1 – Diagram of expression cassettes used in construction of CHROMus mice.** Transgene expression cassettes are placed in frame with the ATG initiation codon from the desired target gene in the desired BAC molecule using standard BAC recombineering protocols. The optogenetic transgene cassettes contain the coding sequence for the optogenetic protein (opto$\alpha$1AR, optoß2AR, or CatCh2) and the bacterial LacZ coding sequence separated by an internal ribosome entry sequence (IRES) element. Expression cassettes for fluorescent reporter proteins contain circularly permutated GFP or RFP coding sequence flanked by synthetic M13 peptide and calmodulin coding sequences, 5' and 3' respectively. In some cassettes, a fusion protein was created by linking an RFP (mCherry or mVermilion) to the carboxy terminus of the GCaMP protein separated by a spacer peptide. The purified recombinant BACs were microinjected into single-cell zygotes at Cornell University or UC-Irvine as described in Materials and Methods.

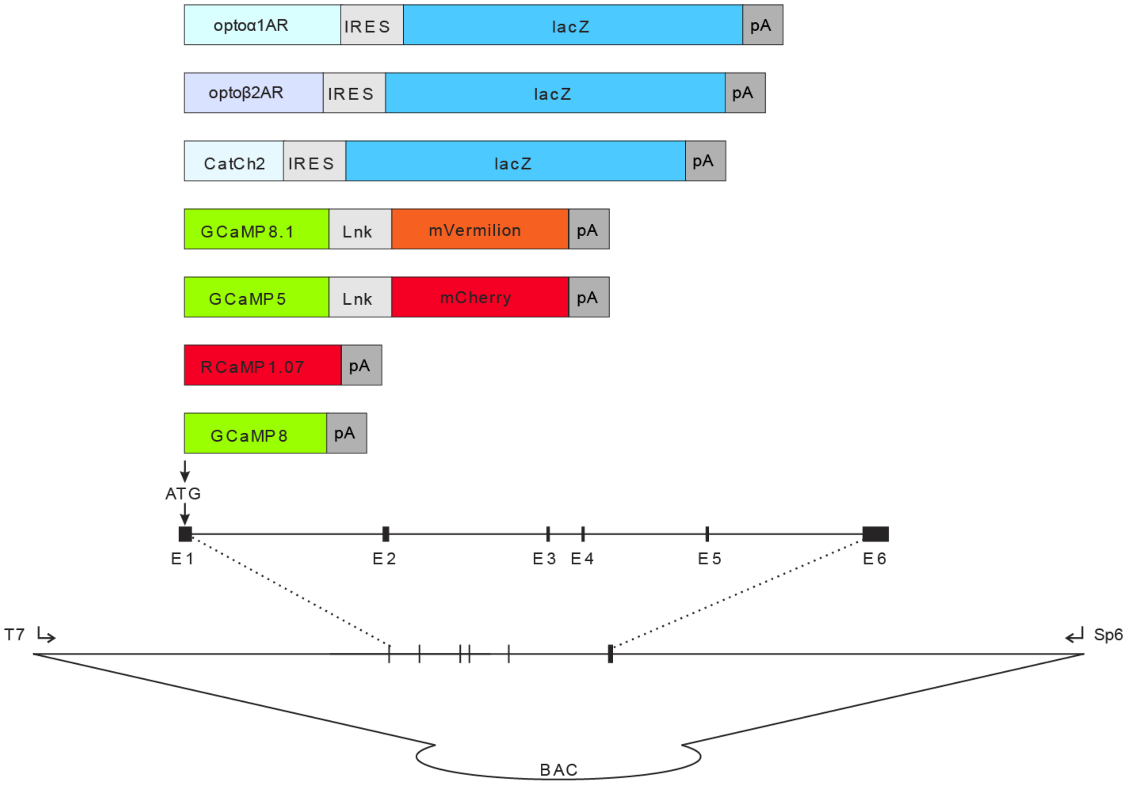

**Figure 2 – figure supplement 1. $HCN4^{BAC}$- CatCh2_IRES_LacZ mice. A**, Ventral view of X-gal stained adult whole heart showing LacZ expressing tissue at the base of the right carotid vein (arrow). **B,** Dorsal view showing LacZ expression along the right carotid vein starting at the junction with right atrium and extending to the base of common carotid vein loop. Ao – aorta, RCV – right carotid vein, RA – right atrium, CCV – common carotid vein. **C,** ECG pacing at 4 Hz can be reestablished after return to native cardiac rhythm. **D,** Laser stimulation of non-CatCh2 expressing tissue does not lead to ECG pacing. All images shown are representative images from 3 animals unless otherwise specified.

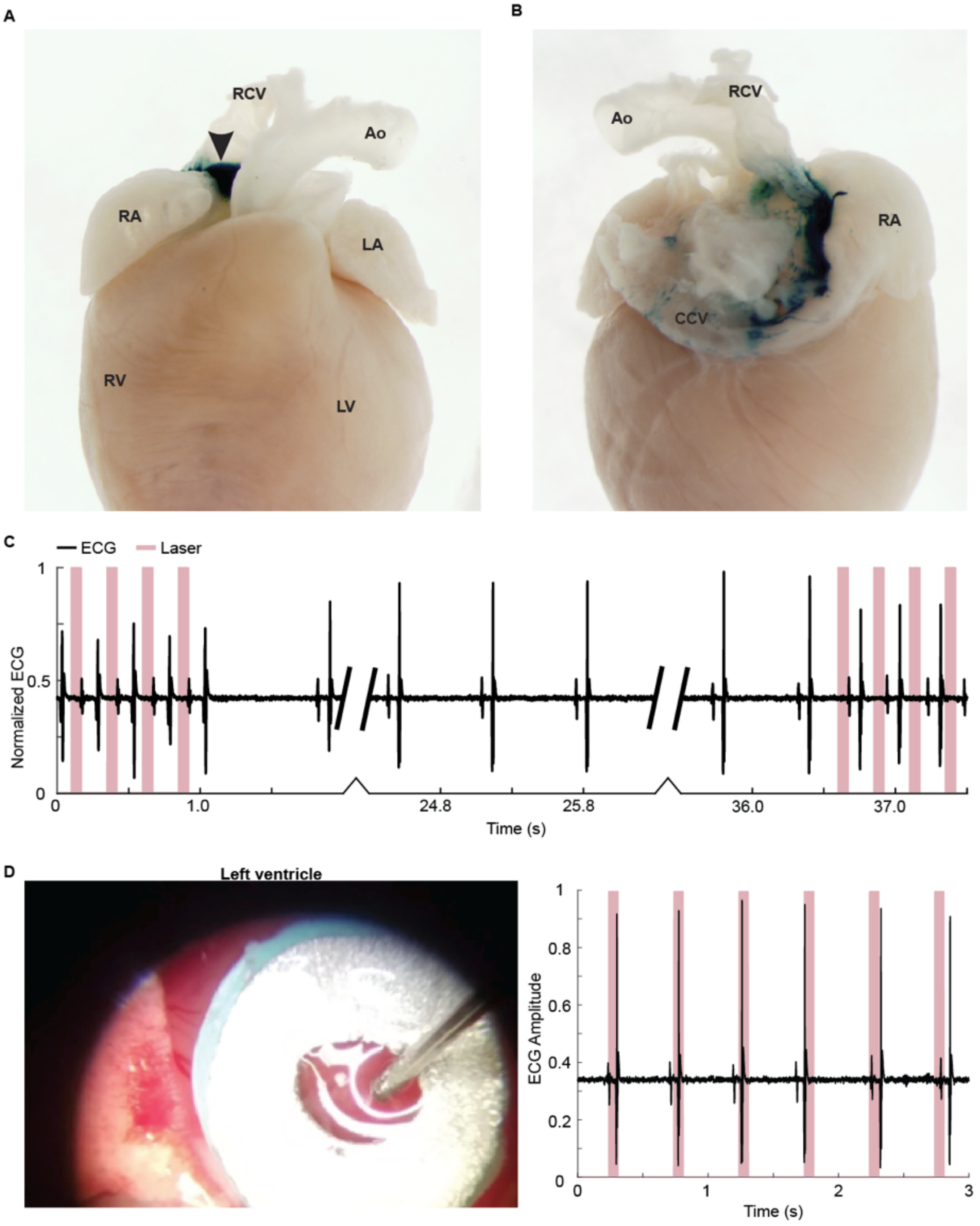

**Figure 2 – figure supplement 2. GCaMP8 fluorescence in hearts from HCN4$^{BAC}$-GCaMP8 mice. A,** Native fluorescence from SA nodal tissue in developing embryos. Left inset: two transgenic embryos and one non-transgenic embryo; arrowhead points to SA nodal tissue with GCaMP8 signal. **B,** GCaMP8 activity in the junction between right atrium (RA) and right ventricle (RV) in a single contraction cycle. The outer wall over the junction was removed to expose the internal tissue. Colorization was achieved using the 16-color LUT in Fiji program and the scale is shown below the images. The box denotes the area of enlargement. All images shown are representative images from 3 animals unless otherwise specified.

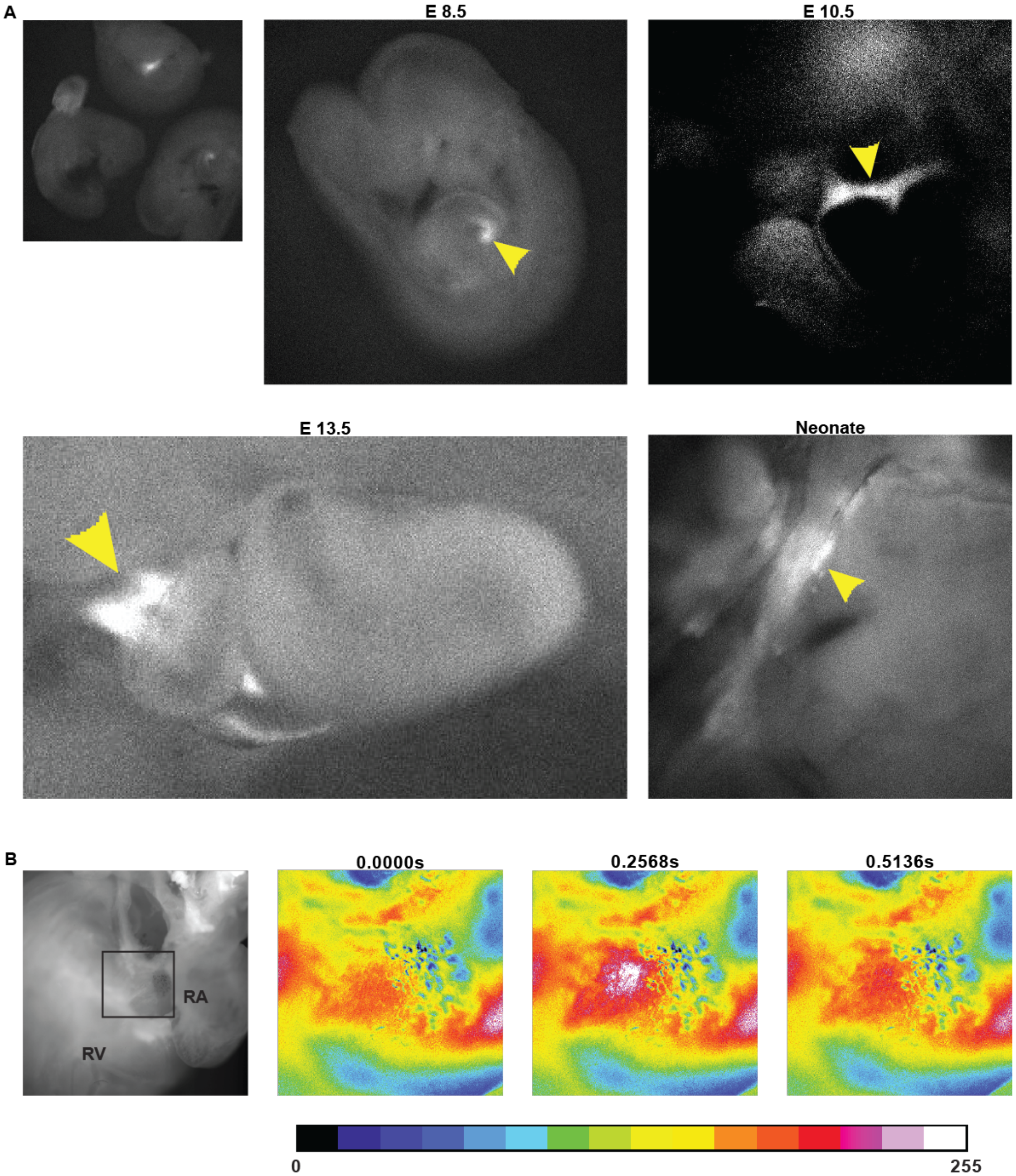

**Figure 4 – figure supplement 1. Acta2$^{BAC}$ optogenetic effector lines. A,** RFP expression in Acta2$^{BAC}$-RCaMP1.07. Upper panel - the RCaMP1.07 is expressed in coronary artery (A) but not veins (V). Scale bar – 100 µm. Lower panel - RCaMP1.07 fluorescence signaling in the right atrium (dashed line – right atrium). **B,** GCaMP5 signal in longitudinal muscle of large intestine (*taeniae coli*) demonstrating intracellular $Ca^{+2}$ concentration changes with muscle contraction and relaxation. Color scale shown below the images. **C,** X-gal staining in various tissues from Acta2$^{BAC}$ effector mice and non-transgenic litter mate. Scale bars; Upper panels – 300 µm, Lower panels – 200 µm. All images shown are representative images from 3 animals unless otherwise specified.

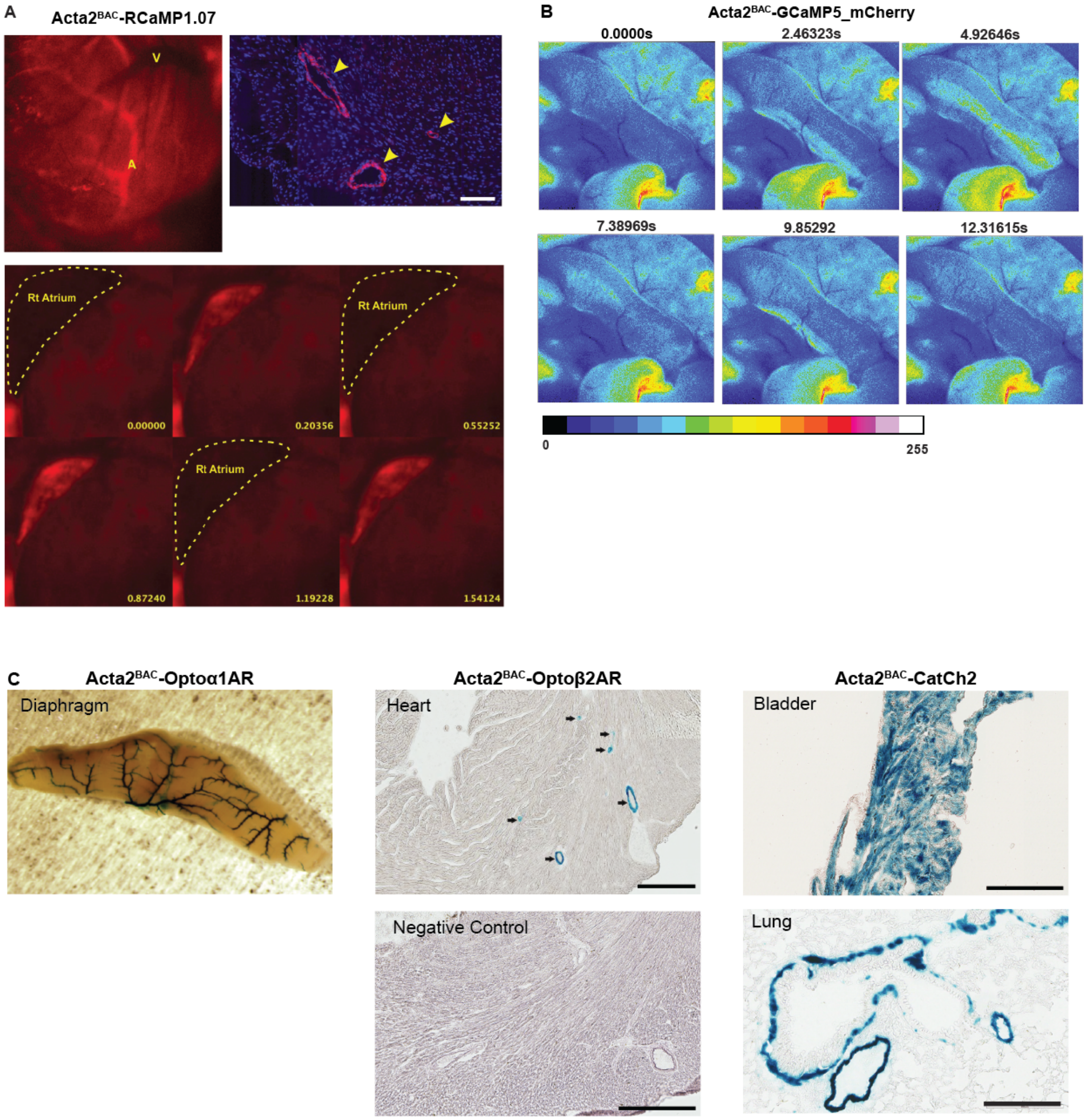

**Figure 6 – figure supplement 1. Cdh5$^{BAC}$ lines. A,** X-gal staining of brain blood vessel from Cdh5 optoeffector mice. **B,** X-gal staining of vasculature in cremaster muscle and lung cryosection from Cdh5$^{BAC}$-Opto$\alpha$1AR mouse. Left panel - cremaster staining; right panel - staining of lung vasculature. Scale bar - 200 µm. **C,** Native fluorescence image from a non-transgenic litter mate shows no GCaMP8 fluorescence compared to transgenic litter mate in Figure 6B. Scale bar – 200 µm. **D,** Laser activation of cremaster artery from Cdh5$^{BAC}$-Opto$\alpha$1AR leads to vessel dilation. All images shown are representative images from 3 animals unless otherwise specified.

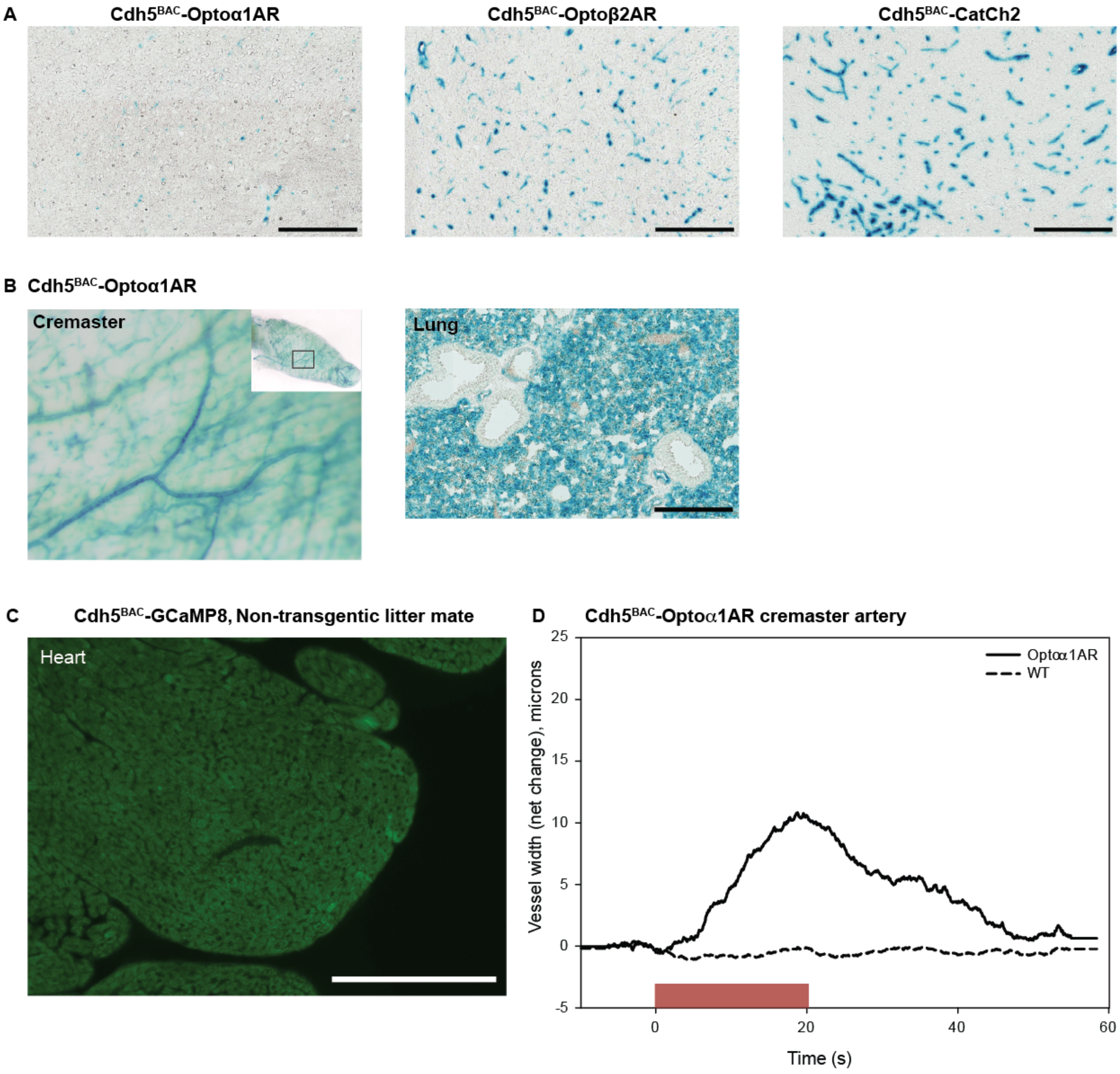

**Figure 8 – figure supplement 1. Transgene expression in other CHROMus lines.** **A,** Anti-GFP IHC of aorta from Cx40$^{BAC}$-GCaMP5_mCherry mouse demonstrates endothelial GCaMP5 expression. Scale bar – 200 µm. **B,** X-gal staining of adrenal gland from a Dbh$^{BAC}$-CatCh2_IRES_lacZ mouse showing LacZ expression in the adrenal medulla. Box indicates area of enlargement. Scale bars; left panel – 500 µm, right panel – 200 µm. **C,** X-gal staining of kidney (left panel) and olfactory lobe (right panels) from Aldh1L1$^{BAC}$-Opto$\alpha$1AR_IRES_lacZ mouse. Box indicates area of enlargement. Scale bars; upper panel – 200 µm, lower panels – 600 and 300 µm. **D,** GCaMP8 expression in lung from SP-C$^{BAC}$-GCaMP8 mice consistent with a Type 2 alveolar cell pattern. Scale bar – 300 µm. All images shown are representative images from 3 animals unless otherwise specified.

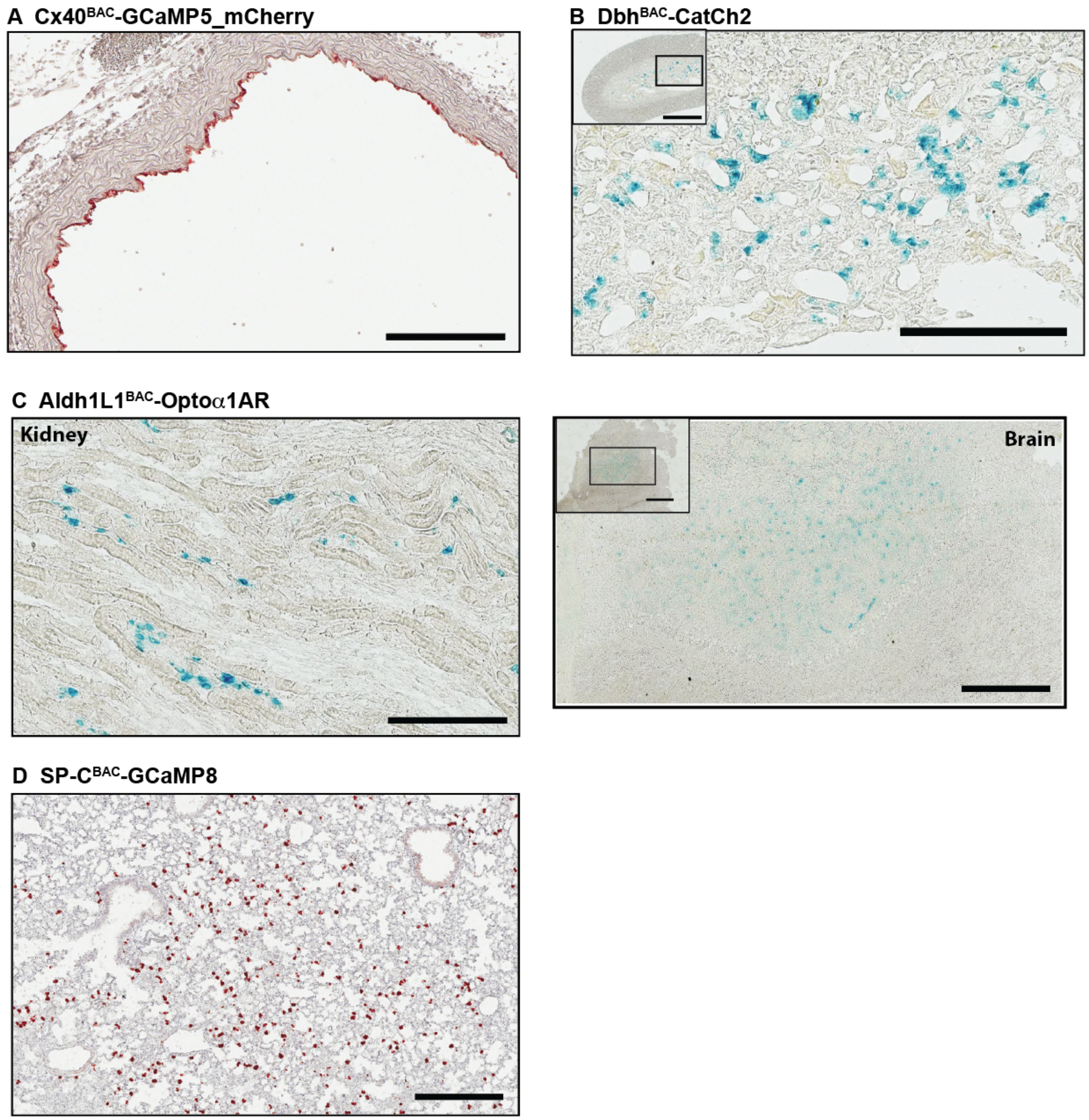

**Supplemental Movies**

**Supplemental Movie 1.** GCaMP8 fluorescence from laser pacing of HCN4$^{BAC}$-CatCh2/αMHC-GCaMP8 heart. Laser stimulation at the SA node in the biallelic heart results in pacing of GCaMP8 fluorescence in the left ventricle. The white strip to the right of the ventricle denotes laser pulse.

**Supplemental Movie 2.** GCaMP8 fluorescence in a E13.5 heart from an HCN4$^{BAC}$-GCaMP8 mouse. GCaMP8 fluorescence migrates from the SA node to the left ventricle.

**Supplemental Movie 3**. GCaMP8 fluorescence in the atrioventricular junction of HCN4$^{BAC}$-GCaMP8 adult heart. The image was colorized in ImageJ, for color scale see Online Figure III.

**Supplemental Movie 4.** Image stack from 2-photon imaging of ventricle contraction in αMHC-GCaMP8. Green- GCaMP8, red – Texas Red dextran in vessels.

**Supplemental Movie 5.** RCaMP1.07 fluorescence in right atrium of Acta2$^{BAC}$-RCaMP1.07

**Supplemental Movie 6.** GCaMP5 fluorescence in large intestine of Acta2$^{BAC}$-GCaMP5_mCherry mouse. The image was colorized in ImageJ, for color scale see Online Figure III.

**Supplemental Movie 7.** Dilation of Acta2$^{BAC}$-Optoß2AR artery stimulated with blue light.

**Supplemental Movie 8.** Vasoconstriction of Acta2$^{BAC}$-CatCh2 artery stimulated with blue light.
